# Tracking the Spread of a Naturally Occurring *Leishmania infantum* Mutant: A qPCR-Based Investigation in Strains and Clinical Samples

**DOI:** 10.1101/2025.08.22.671732

**Authors:** Marne Coimbra das Chagas, Ana Nilce Silveira Elkhoury, Samantha Valadas, Bruna Dias das Chagas, Camilly Enes, Artur Augusto Velho Mendes Junior, Sandro Antonio Pereira, Gabriel Carvalho de Macedo, André Luiz Rodrigues Roque, Manuela Solcà, Claudia Brodskyn, Deborah Bittencourt Mothé Fraga, Juliana Mariotti Guerra, José Eduardo Tolezano, Luiz Fabio Batista, Marcia Dalastra Laurenti, Lisvane Paes-Vieira, Carlos Robello, Otacilio C. Moreira, Elisa Cupolillo, Mariana C. Boité

## Abstract

The circulation of *Leishmania infantum* strains carrying a 12 Kb genomic deletion (DEL) alongside wild-type (NonDEL) parasites in the Americas raises critical epidemiological questions. To support molecular surveillance efforts, we developed and validated a qPCR-based genotyping tool capable of detecting and quantifying DEL, NonDEL, and coinfection (MIX) profiles. The strategy targets a constitutive region present in all strains and a specific region exclusive to NonDEL parasites. It was validated using DNA from cultured parasites and applied to a panel of diverse clinical samples from dogs, humans, and other hosts across South and Central America. DEL genotypes were found to be more frequent and widespread, both in clinical samples and cultured parasites, suggesting their significant role in the transmission cycle. The tool’s application revealed coinfections in paired samples and enabled relative quantification of genotypes. It also showed good correlation with parasite load estimations and high discriminatory power when integrated with the Genotype function of real-time PCR software. This protocol provides a robust, scalable approach for monitoring *L. infantum* genotypes, with direct implications for understanding infection dynamics, treatment outcomes and public health surveillance.

**Author Summary:** *Leishmania infantum* is the parasite responsible for visceral leishmaniasis in the Americas. Recent studies have shown that two distinct genotypes—those carrying a specific gene deletion (DEL) and those without it (NonDEL)—coexist and circulate in the region. Understanding how these genotypes are distributed and whether they affect disease transmission or treatment is critical, especially because Miltefosine, a drug used to treat dogs, may not work equally well against both genotypes. In this study, we developed a simple and accurate molecular test to detect and quantify these genotypes in both cultured parasites and clinical samples from different animal hosts, including humans. We found that DEL genotypes are common and widely distributed, and that some animals carry both types of parasites simultaneously. Our method also distinguishes *L. infantum* from other related species, making it useful for diagnosis and surveillance. This tool can help researchers and health authorities monitor parasite populations more effectively and better understand how different genotypes might impact disease outcomes and control strategies.

## Introduction

*Leishmania infantum* is the agent of American Visceral Leishmaniasis (AVL), a lethal disease if left untreated. The main urban reservoir is the domestic dog, which also can develop the disease, leading to Canine Visceral Leishmaniasis (CVL). The transmission cycle includes vector sandflies of various species, depending on the region. *Lutzomyia evansi* was described as participating in the transmission of the parasite in Colombia and Venezuela(1,2). The main vector is, however, *Lutzomyia longipalpis*(3). *Lutzomyia cruzi* is found in municipalities in the border areas between Bolivia and Brazil and between Bolivia and Argentina(2). *Lutzomyia migonei* has also been described as a competent vector for *L. infantum*(4) in Brazil and in Argentina(5).

Epidemiological changes of AVL are noteworthy and, mainly, the occurrence of the disease continues to expand geographically, in both humans and dogs(6,7). The overall number of cases in the Americas has diminished in the recent years, but when countries are analyzed separately, Brazil harbors more than 90% of human AVL cases(8), and Paraguay and Argentina reveal an increase in the numbers(2). Even though the absolute number of deaths lowered in the Americas, the lethality rate increased(8). These modifications may be related to a complex overlapping of different factors. The adaptation of the vector sandfly species to the urban environment, for instance, is quite relevant in the disease expansion scenario.

Another major player that may lead to modifications on the AVL scenario is the parasite itself. Studies have demonstrated that the parasites that circulate in the Americas belong to the same species present in Europe, Africa, and Asia, that arrived in the Americas during the last five centuries(9,10). The consequent bottleneck effect during such parasite importation was the adaptation to novel vertebrate (native fauna) and invertebrate hosts (vector species that are not present in the Old World), possibly fostering genomic variation over the New World *L. infantum* population, which has been under investigation by researchers. Although frequently considered a homogeneous population, the American strains of *L. infantum* (syn. *L. chagasi*) have been demonstrated to correspond at least to three distinct populations(11) by multilocus microsatellite analysis (MLMT). As a consequence of investigation efforts, including comparative deep sequencing analysis, a specific and relevant genomic finding has been published: a 12 KB sub-chromosomal deletion, described exclusively among Brazilian strains of *L. infantum* (12,13).

Some epidemiologically relevant phenotypes correlate with this parasite’s genomic trait. Our group recently described higher metacyclogenesis in vitro and in vivo for deletion-carrying parasites (DEL), and an ability to infect the vertebrate host despite its susceptibility to the first cell line defense(14). These findings suggest that DEL mutant strains could alter pathogenicity and lead to less symptomatic clinical outcome. Emphasizing the main urban reservoir, the domestic dog, this might represent a threat to surveillance and disease control since these animals could remain for longer periods undetected as a source of infection for the vector, contributing to keeping the endemicity in certain areas.

Notably, the deletion has been associated with reduced susceptibility to miltefosine (MIL)(12,13), one important drug for treating the human disease in several countries outside the Americas, but approved to treat CVL only in Brazil. This fact raises concerns for surveillance and control, because the parasite population circulating is the same among humans and dogs. The dogs are often asymptomatic and, when symptomatic, can present a wide spectrum of clinical manifestations, irrespective of their parasite load. The concern is that the treatment focuses on the clinical recovery of the dogs, a goal indeed achievable(15); but it is still unclear how infective to vectors these dogs may remain(16), and what would be their role in maintaining the presence of the parasite in urban centers. Therefore, monitoring the outcome of infected / treated dogs by measuring parasite load may contribute to these investigations. The alarm is magnified by the fact that the less susceptible strains to MIL, i.e., the DEL parasites are also the more frequent and widespread in Brazil, as reported previously by our group(17). In that study the authors reported that Non-deleted strains (NonDEL) appeared concentrated in a few Brazilian states, and in areas with co-circulation of both DEL and NonDEL, heterozygous strains were also detected(17), suggesting co-infections by both genotypes occur.

In this scenario, the parasite’s genomic site of the deletion may represent a pivotal molecular marker for clinicians and surveillance to follow up the epidemiology of VL in the country. However, the currently known distribution of the *L. infantum* DEL and NonDEL genotypes is based exclusively on isolated parasites (convenient samples) kept in a biological collection. Based on these statements, the goal of the present work is to present a detailed mapping of the occurrence of genotypes in clinical samples, avoiding the potential bias of culture-selection. To achieve this aim, a Real time qPCR-based approach was applied to detect, quantify and to type *L. infantum* as DEL and NonDEL in clinical material from different sources and hosts. The obtained results updated the distribution and relative frequency of the DEL and NonDel genotypes along the Americas, offering an accurate and reliable qPCR-based approach able to quantify DEL and NonDEL parasites in clinical material, even in mixed infections.

## Methods

### Samples and DNA preparation

Parasites were donated by the Coleção de *Leishmania* da FIOCRUZ (CLIOC - https://clioc.fiocruz.br). The *Leishmania infantum* strains were previously characterized as DEL or NonDEL in a former study, either by qPCR or by whole genome sequencing(14,17). The strains were kept in culture and prepared for further DNA isolation by the High Pure PCR Template Preparation Kit (Roche) in accordance with the manufacturer protocol. The species was confirmed either by multilocus enzyme electrophoresis (MLEE), also in accordance with internal procedures or by PCR and DNA sequencing. Data was gathered from 238 different strains.

DNA from clinical samples was obtained from the Reference service and from studies and collaborators from Brazil and other American countries. Samples could be either replicates from the same host or single samples. In total, 292 different individuals were included, and the panel was composed of samples from dogs (n=369), cats (n=2), coatis (n=10) and phlebotomines (n=3). The clinical material was fragments of healthy skin, skin lesions, bone marrow aspirate, lymph node aspirate and spleen, and, additionally, the intestines from the phlebotomines (Supp. Material). The material was shipped to the Laboratório de Referência e Diagnóstico para *Leishmania* (LRDIL) to confirm the presence of *Leishmania* DNA and for the typing of the infecting species by molecular methods, i.e., conventional PCR targeting the ITS1 region of *Leishmania* sp (18) followed by sequencing.

### Ethical statements

All samples used represent leftover biological material collected for the clinical care of dogs and other studies. This activity falls under option b-ii of item 6.1.10 of the annex to Normative Resolution No. 55 (CONCEA, October 2022) and therefore does not require an ethical license. A Declaration Letter of the Coordinator of AEC/Oswaldo Cruz Institute stating that the project does not require an ethical license is available upon request.

### qPCR design and validation of the protocol

Primers and probes were designed targeting distinct regions of Chromosome 31, one of which the deletion is located(12). The available nucleotide sequence for *Leishmania infantum* JPCM5 genome - chromosome 31 was used as a reference, combined with other available genomes, for selecting conserved regions of the *L. infantum* genome. To cover the deleted sub-chromosomic region, the NCBI Reference Sequence NC_009415.2 was used as a template in Primer Blast (https://www.ncbi.nlm.nih.gov/tools/primer-blas). As a constitutive target, we selected a downstream region within chromosome 31 existent in both, DEL and NonDEL *L. infantum* strains. The region (NCBI Reference Sequence: XM_001467491.1) codes for metallo- peptidase, Clan MA(E), Family M1. The nucleotide sequences for primers and probes are depicted in Table 1. Available nucleotide sequences for the hypoxanthine phosphoribosyltransferase 1 gene (HPRT) from domestic dog (*Canis lupus familiaris*) were used as template and as an endogenous target for the quantitative approach (Table 1). For clinical material from other hosts, the 18s SSU target conserved for all mammals was applied in conventional PCR and endogenous control for the routine workflow (Primers: F: 5’ CGG CTA CCA CAT CCA AGG AA 3’; R: 5’ CCT GTA TTG TTA TTT TTC GTC ACT ACC T 3’).

**Table 1.**
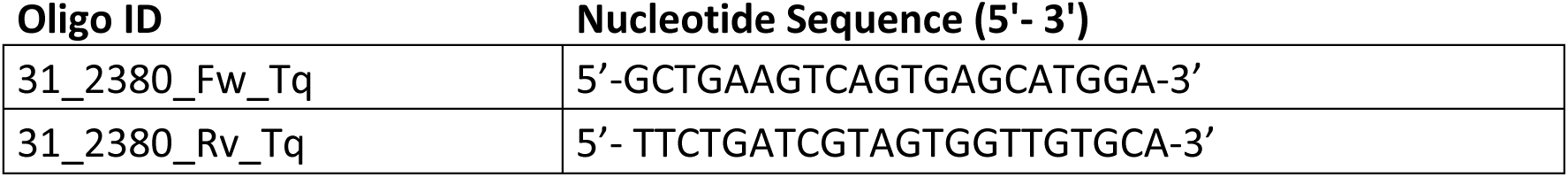
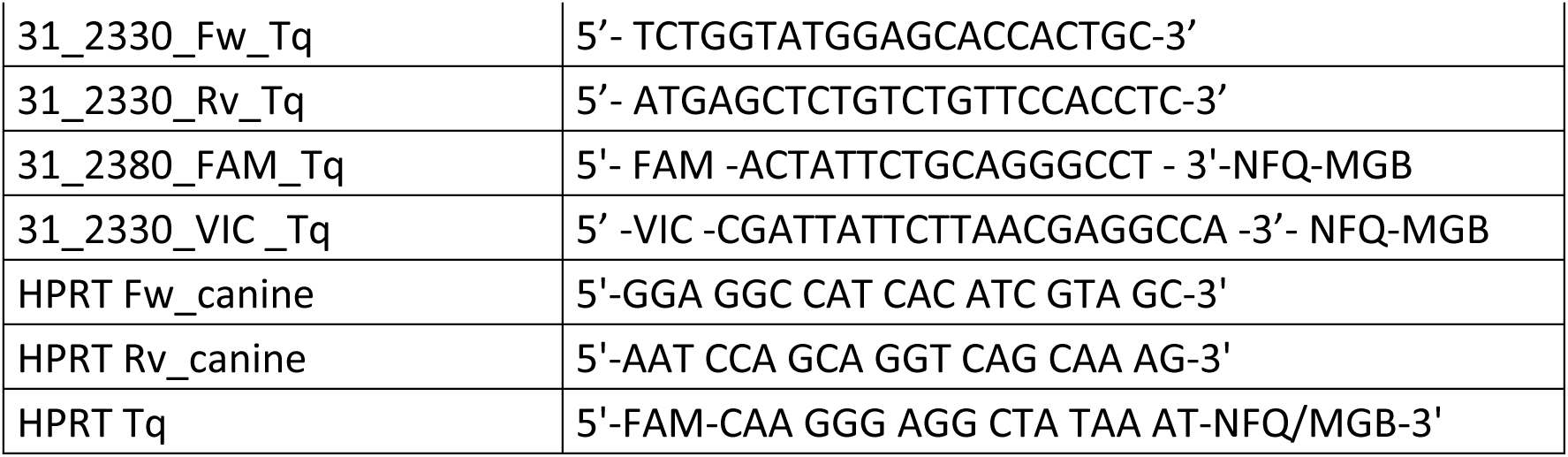
Primers and probes chosen based on primer Blast results.

Final concentration of primers and probes was obtained after assays testing a gradient of three concentrations. The protocol includes the TaqMan™ system (Applied Biosystems® CA, USA) and the protocol was defined as 0.15 µM of each primer and 0.100 µM of probes. The assay was performed, according to the following conditions: 45 cycles of 95° 15 sec, 61 ° 1min. qPCRs were carried out with Applied Biosystems VIA7 equipment and Quant Studio Real-Time from the Plataforma de Análises Moleculares (RPT09J) (Rede de Plataformas Tecnológicas Fiocruz). A DNA-free master mix reaction (*No Template Control* - NTC) and all samples and controls were performed in duplicate.

### Genotyping setting

Genotyping experiments in real time qPCR can be designed to detect single nucleotide polymorphisms (SNP) variant of a target nucleic acid sequence. This set up thus allows discrimination between alleles, representing an unbiased approach to differentiate DEL vs NonDEL *L. infantum* and other *Leishmania* species. To test the accuracy of the Genotype qPCR setting, available within the QuantStudio3 software (Design and Analysis 2 – DA2 v2.8.0, Applied Biosystems) a panel of DNAs from representative reference strains from different *Leishmania* species (representing *Leishmania (Viannia)* sp., *L. (Leishmania) sp., L. (Porcisia)* and Paraleishmania) was prepared for the multiplexed genotyping assay. In total, 14 different species were represented. Samples belonging to the *L. donovani* complex from the Old World (OW; n=3) and from Brazil, representing DEL (n=5), HTZ (n=1), and noNDEL (n=5) genotypes were included. Multiplexed reactions with both targets were prepared as depicted in Table 2.

**Table 2.** Reaction details for the multiplexed protocol prepared for the Genotyping setting.

| Multiplex 31_2330 e 31_2380 |  |  |  |  |  |
| --- | --- | --- | --- | --- | --- |
| Reagent | Concentration |  | Concentration/<br>reaction |  | Volume/<br>reaction µL |
| Water | - | - | - | - | 6,4 |
| Master Mix |  |  |  |  |  |
| Taqman | 2 | x | 1 | x | 10 |
| 31_2330_Fw_Tq | 10 | microM | 0,15 | microM | 0,3 |
| 31_2330_Rv_Tq | 10 | microM | 0,15 | microM | 0,3 |
| 31_2330_VIC_Tq | 10 | microM | 0,1 | microM | 0,2 |
| 31_2380_Fw_Tq | 10 | microM | 0,15 | microM | 0,3 |
| 31_2380_Rv_Tq | 10 | microM | 0,15 | microM | 0,3 |
| 31_2380_FAM_Tq | 10 | microM | 0,1 | microM | 0,2 |
| DNA | - | - | - | - | 2 |

### Mapping the distribution of genotypes

The mapping representation was obtained using the *QGIS Geographic Information System. Open Source Geospatial Foundation Project* (19) - http://qgis.osgeo.org). The input was prepared by the conversion of information of samples origins (locality) into geolocation data available for the municipality (Supp material). The frequencies of detection of DEL and NonDEL for each covered region were added in a table format used as in input for QGIS. The software then generates an output of the mapping of genotypes and the frequencies in a pie chart model.

### Determination of parasite load

Standard curves were obtained by the addition of 10^8^ parasites into 20 mg of DNA derived from skin samples collected from non-infected dogs. The DNA of the material was extracted with the High Pure PCR Template Preparation Kit (Roche) in accordance with the manufacturer protocol. The DNA was diluted 1:10 in series of seven (starting from 10^8^ parasites into 20 mg), and included in the qPCR for each target (2330, 2380 and HPRT; and kDNA)

### Blend of genotypes and differential quantification

The NonDEL IOCL2666 and the DEL IOCL3598 strains were selected to the controlled blend of DEL and NonDEL parasites for further genotype-specific qPCR quantitation. This assay was necessary for differential quantification in coinfected samples. Parasites were cultivated in Schneider with 20% Fetal Calf Serum and 2% filtered urine at °26 C. After 72 hours of culture from an initial inoculum of 10^6^ cells, the parasites were centrifuged at 4.000RPM and resuspended in 1ml of PBS1x. The content was counted in a Neubauer chamber under 20x microscopy. Nine 1.5ml tubes were prepared, each containing 10^7^ cells of the DEL strain. To the first tube we added 10^7^ NonDEL cells and a serial dilution prepared on the remaining 8 tubes already containing 10^7^ DEL cells. The nine tubes thus presented a blend with fixed amount of DEL cells and a serial dilution of NonDEL content. The DNA of each tube was extracted by the High Pure PCR Template Preparation Kit (Roche) in accordance with the manufacturer’s protocol and submitted to the qPCR. The quantity obtained by qPCR was plotted against the quantity obtained by counting (microscopy).

## Results

### Genotypes were accurately detected by the protocol in various types of material, revealing that DEL parasites are more frequent and geographically spread in the Americas despite the origin of the material

DNA from culture parasites representing a panel of *L. infantum* strains previously characterized as DEL and NonDEL by whole genome sequencing (WGS)(17) was used to validate the protocol (Table 3). Results were interpreted after the confirmation of the amplification of the *Leishmania*-constitutive region 31_2330 in all samples. Genotype was defined as NonDEL if 31_2380 also amplified (present) and, as DEL if the 2380 target did not amplify. Figure 1 depicts the strategy of using both targets to offer a typing diagnostic tool for infection by both *Leishmania infantum* genotypes. Primers and probes were named accordingly, i.e., 31_2330 for the constitutive site and 31_2380 for the deleted region.

**Figure 1.**
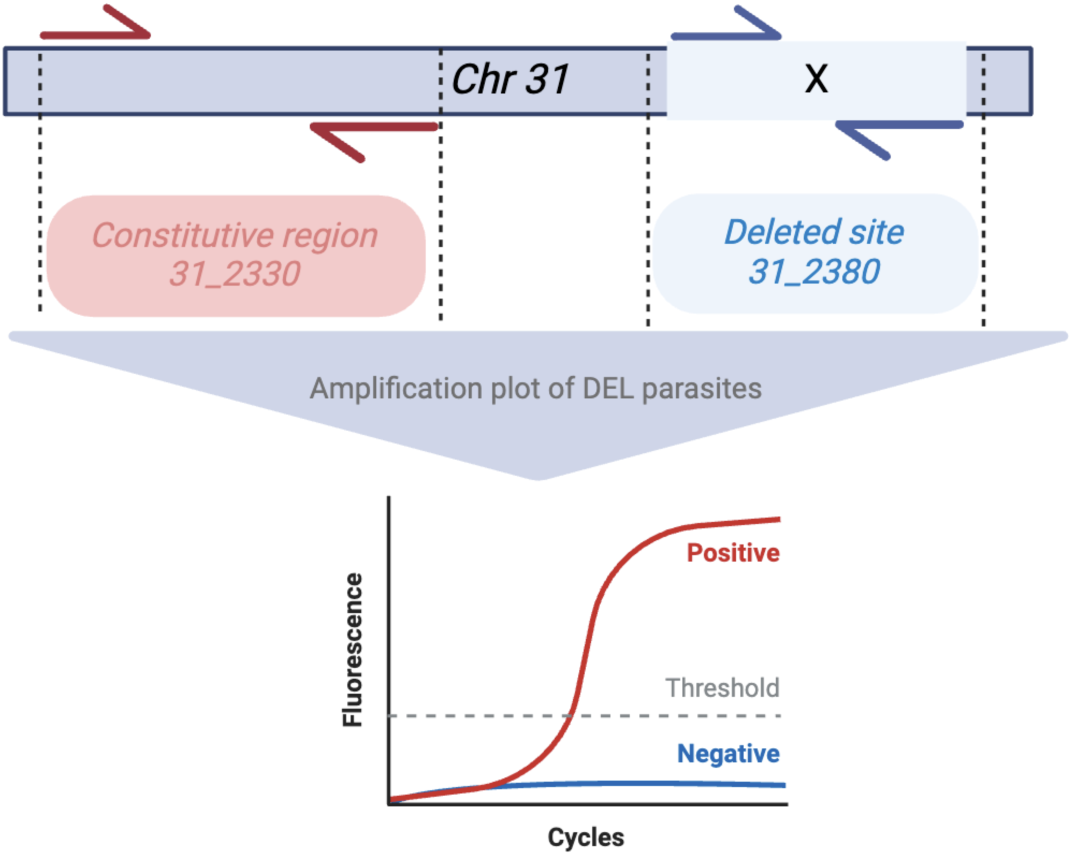
Schematic representation of primers and probes designed to differentially detect L. infantum carrying the deleted site on chromosome 31. In red in represented the primers designed for the constitutive region, present in both DEL and NonDEL samples. The blue area represents the deleted site, amplified uniquely for NonDEL strains. At the bottom is the schematic amplification plot obtained for DEL samples.

**Table 3.** Strains selected to validate the protocol. Collection voucher, international code, genotype by WGS and confirmed by the current protocol, geographic origin and host. The parasites were obtained from CLIOC, and DNA was isolated from cultures using QIAGEN isolation kit. Species was confirmed by Multilocus Enzyme Electrophoresis (MLEE) and sequencing. PE=Pernambuco; PI= Piaui; SE= Sergipe; MT=Mato Grosso; RJ= Rio de Janeiro; SC= Santa Catarina; SP=São Paulo; MS=Mato Grosso do Sul; BA= Bahia; MA=Maranhão.

| Voucher CLIOC | International code | Genotype | Geographic origin | Host |
| --- | --- | --- | --- | --- |
| 2769 | MCAN/BR/2005/SPACK | DEL | PE | canine |
| 2976 | MHOM/BR/2006/1406MBS | DEL | PI | Human |
| 3116 | MHOM/BR/2009/HU-UFS11 | DEL | SE | Human |
| 3223 | MCAN/BR/2010/BOLINHA-III | DEL | MT | canine |
| 3227 | MCAN/BR/2010/MAGRÃO | DEL | MT | canine |
| 3598 | MCAN/BR/2015/TUBO9 | DEL | RJ | canine |
| 3684 | MHOM/BR/2006/ARS 3P | DEL | PI | Human |
| 3729 | MCAN/BR/2016/PRETO | DEL | SC | canine |
| 3634 | MCAN/BR/2016/90 | DEL | RJ | canine |
| 3134 | MCAN/BR/2009/GRANDÃO-I | HTZ | MT | canine |
| 2949 | MCAN/BR/2004/LIBPI-18 | HTZ | PI | canine |
| 3254 | MCAN/BR/2009/CLV17 | HTZ | SP | canine |
| 2666 | MCAN/BR/2002/LVV-137 | NonDEL | MS | canine |
| 2848 | MCAN/BR/2005/NMT-BSB159B | NonDEL | BA | canine |
| 2972 | MHOM/BR/2005/742EMS | NonDEL | PI | Human |
| 3124 | MHOM/PT/1989/IMT-202 | NonDEL | Portugal | Human |
| 3210 | MCAN/BR/2010/CALSITO-I | NonDEL | MT | canine |
| 3556 | MCAN/BR/2014/18MEDULA | NonDEL | MA | canine |
| 3685 | MHOM/BR/2006/MA03-A 4P (GMS) | NonDEL | PI | Human |
| 2668 | MCAN/BR/2002/LVV-139 | NonDEL | MS | canine |
| 2584 | MHOM/BR/2003/JBIC | NonDEL | MS | Human |
| 2664 | MCAN/BR/2002/LVV-135 | NonDEL | MS | canine |
| 3737 | MCAN/BR/2017/BETHOVEN | NonDEL | SC | canine |

The outcome revealed 100% accuracy for detecting the genotypes ( revealing the non-equivalent quantity of targets. This trait allows the protocol to be applied as a genotype-specific quantitative tool.

Figure 2). Moreover, amplification plots of NonDEL samples show similar Ct values for both targets (+/- 0.5) ( revealing the non-equivalent quantity of targets. This trait allows the protocol to be applied as a genotype-specific quantitative tool.

**Figure 2.**
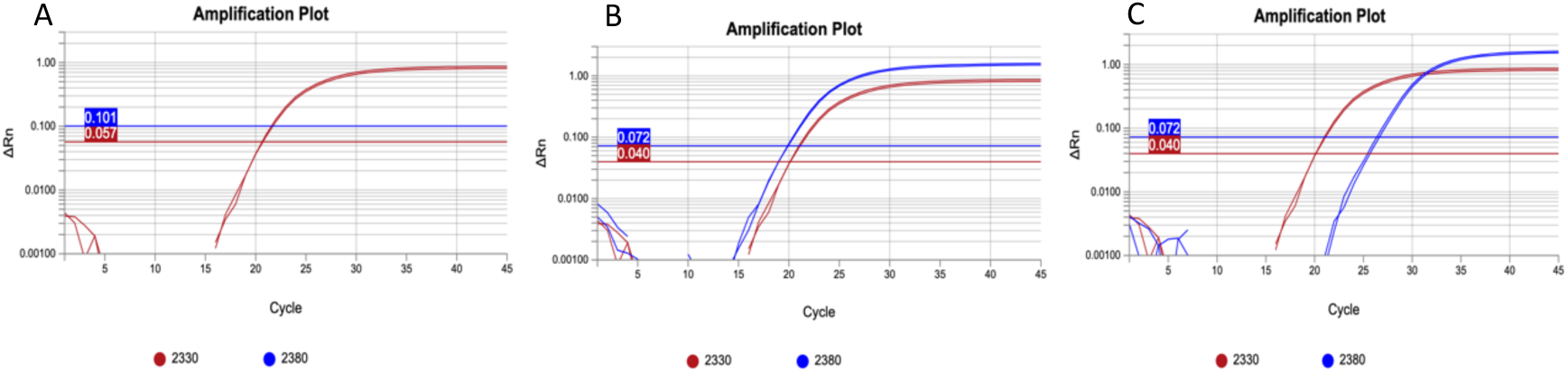
qPCR amplification plot of the constitutive target 31_2330 and the deleted site 31_2380 in reference samples for DEL, NOnDEL and HTZ genotypes. A) DEL sample (IOCL3598) showing only the amplification of 2330 (constitutive site). B) NonDEL sample (IOCL 2666) showing both amplifications with similar Cts values. C) HTZ sample (IOCL 3134) showing the deviation between Cts – 2380 presents lesser number of copies due to the presence of the deletion only in half of the homologue chromosomes.

Figure 2B). The detection of samples infected with either heterozygous parasites or co-infection by DEL and NonDEL can be done by the detection of Cts deviation (deltaCt > 0.5) between targets ( revealing the non-equivalent quantity of targets. This trait allows the protocol to be applied as a genotype-specific quantitative tool.

Figure 2C) revealing the non-equivalent quantity of targets. This trait allows the protocol to be applied as a genotype-specific quantitative tool.

After confirming the protocol as a typing tool for *L. infantum* genotypes a wider panel of samples (n=622) was prepared including DNA from cultured parasites (n=238) and clinical samples (n=384) from different hosts and tissues. Most clinical samples in the present study are from dogs (n=369), which are more accessible and a valuable material for surveillance. Therefore, specific primers and probes were also designed as internal constitutive targets for canine hosts’ DNA (HPRT; Methods), for use as sample-quality control and for parasite load. The criterion for defining the results is depicted in Table 4, considering the profiles represented in Figure 2, as follows: DEL = 31_2330 amplified, 31_2380 negative. NonDEL = both 31_2330 and 31_2380 amplified with overlapped Cts values (+/-0.5). Mix/Htz = both 31_2330 and 31_2380 amplified with deviating Cts values > 0.5. Inconclusive refer to those samples among which amplification was detected for 31_2330 and/or 31_2380 with Cts>35. Negative samples = no amplification detected. For the clinical samples, the protocol presented a percentage of positivity of 90.2% compared to the conventional PCR and allowed 80.6% of assayed clinical samples to be typed as DEL, NonDEL and MIX/Htz.

**Table 4.** Number of samples assayed for the presented protocol and outcomes. All clinical samples were previously confirmed by conventional PCR for the target ITS1(18) or kDNA. DEL = 31_2330 amplified; 31_2380 negative. NonDEL = 31_2330 and 31_2380 amplified with almost overlapped Cts values (+/-0.5). Mix/Htz = 31_2330 and 31_2380 amplified with Cts values deviating > 0.5. Inconclusive = refer to those samples among which amplification was detected for 31_2330 and/or 31_2380 with Cts>35. Negative samples = no amplification detected.

|  | DEL | NonDEL | Mix/HTZ | Inconclusive | Negative | TOTAL |
| --- | --- | --- | --- | --- | --- | --- |
| **clinical samples** | 192 (50.0%) | 69 (18.0%) | 19 (4.9%) | 68 (17.7%) | 36 (9.4%) | 384 |
| **Strains** | 170 (71.4%) | 52 (21.8%) | 16 (6.7%) | 0 | 0 | 238 |
| **TOTAL** | 362 (58.2%) | 121 (19.5%) | 35 (5.6%) | 68 (10.9%) | 38 (5.8%) | 622 |

To map the distribution of genotypes and frequencies the inconclusive and negative samples were excluded. Georeferenced localities (n=57), mainly from Brazil, but also from Bolivia, Uruguay, Honduras and Panama were represented by the positive and successfully typed clinical samples and strains (Figure 3). The deleted genotype appeared more frequently (70%) and spread. Regions such as São Paulo, Mato Grosso, Mato Grosso do Sul and Piaui present more evidently the co-circulation of DEL and NonDEL. Moreover, samples from these regions commonly present a mixed profile, suggesting the coinfection by both genotypes or the presence of heterozygous parasite, as depicted previously (Figure 2C).

**Figure 3.**
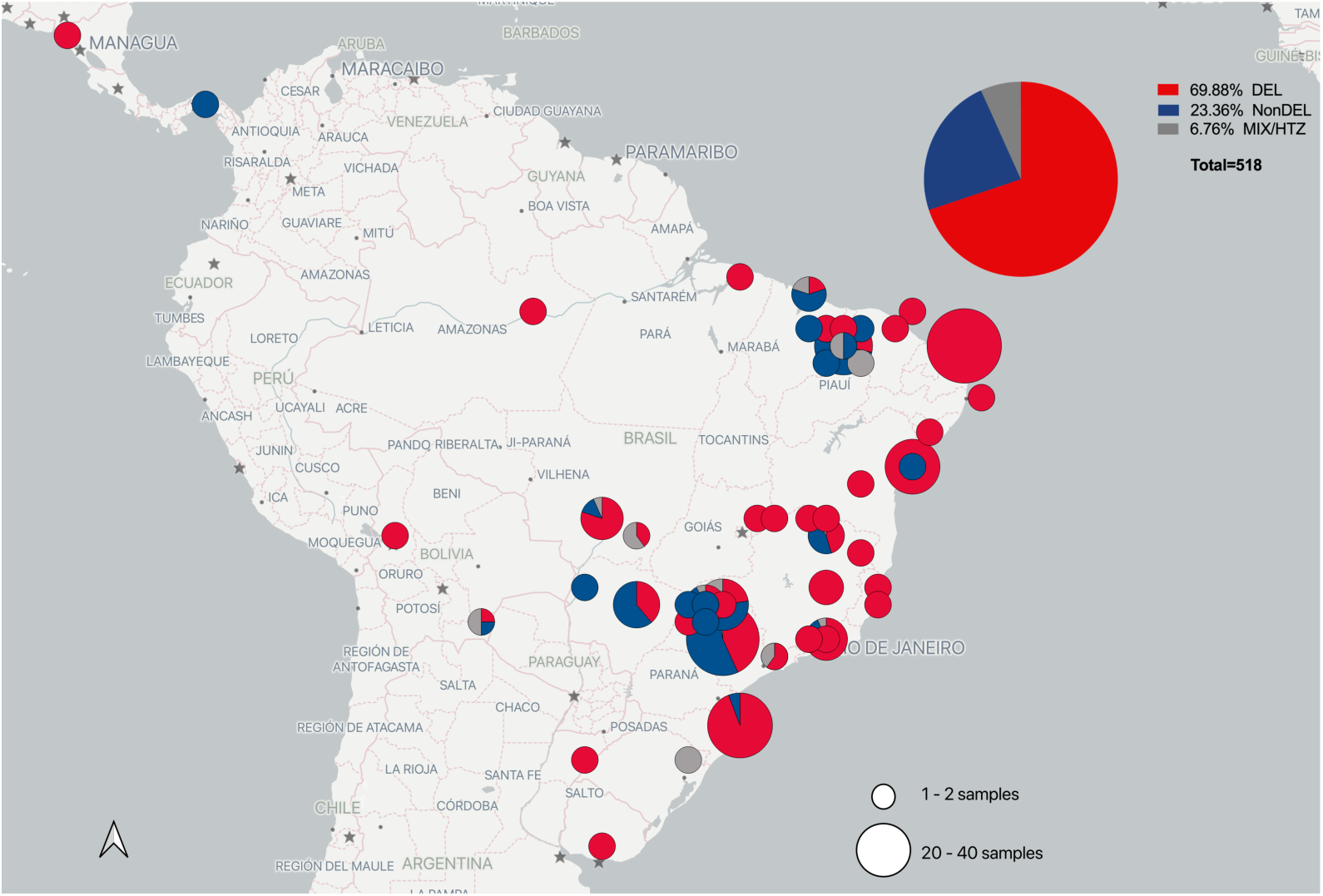
Map covering part of Central and South America showing the relative frequency and geographic distribution of genotypes. The panel of samples comprises both strains (n=238) from CLIOC and clinical material (n=280). Inconclusive and negative samples from Table 3 were excluded and the remaining 518 samples (clinical and strains) mapped accordingly to the georeferenced locations from collecting data. The percentages reflect the higher frequency of DEL samples (depicted in the pie chart on the upper right). Figure with map and frequencies obtained by QGIS (19).

### The Genotyping qPCR setting differentially detected *L. donovani* complex species and identified DEL and NonDEL parasites in clinical samples

To test the accuracy of the method, a panel of DNAs from representative reference strains from different *Leishmania* species (representing *Leishmania (Viannia)* sp., *L. (Leishmania) sp., L. (Porcisia),* and Paraleishmania) was prepared for the multiplexed genotyping assay (Table 5). The software generates an Allelic Discrimination Plot, depicting the fluorescence detected by each probe (31_2330 and 31_2380) and clustering samples accordingly to the amplification efficiency. The clusters are color-coded and corresponds to color-coded results are depicted in Table 5.

**Table 5.**
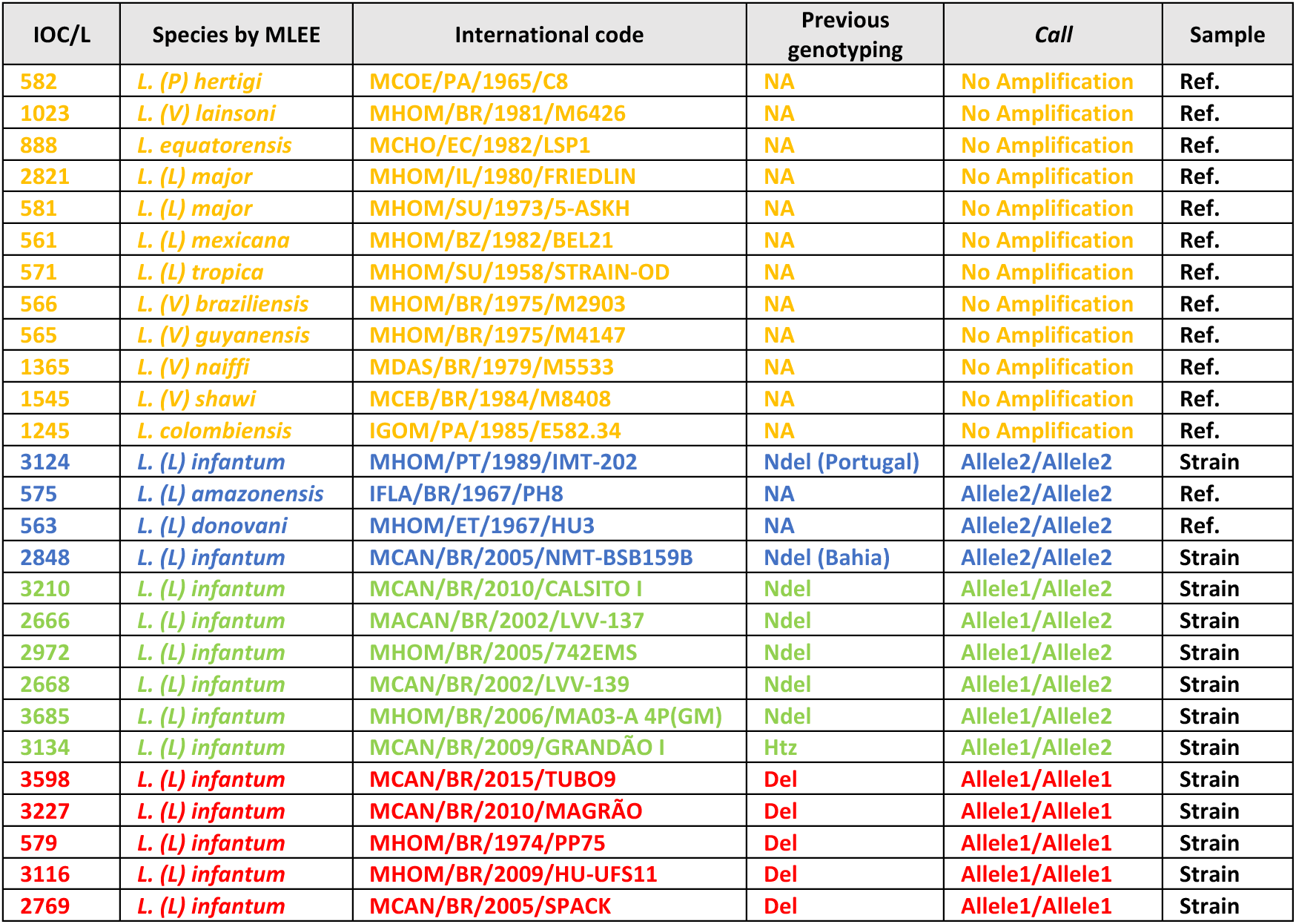
Samples selected to test the differential genotyping approach by the Real time qPCR. Representative samples from the Del and NonDEL genotypes and reference strains from subgenus L. (Viannia), L. (Leishmania), L. (porcisia) and Paraleishmania were included to cover a broad range of Leishmania sp. diversity. IOC/L (voucher code); species determined by Multiloculs Enzyme Electrophoresiss (MLEE); International code; genotyping determined previously by WGS and /or qPCR; “Call” represents the color-coded clustering determined by the software; type of sample.

| IOC/L | Species by MLEE | International code | Previous genotyping | Call | Sample |
| --- | --- | --- | --- | --- | --- |
| 582 | *L. (P) hertigi* | MCOE/PA/1965/C8 | NA | No Amplification | Ref. |
| 1023 | *L. (V) lainsoni* | MHOM/BR/1981/M6426 | NA | No Amplification | Ref. |
| 888 | *L. equatorensis* | MCHO/EC/1982/LSP1 | NA | No Amplification | Ref. |
| 2821 | *L. (L) major* | MHOM/IL/1980/FRIEDLIN | NA | No Amplification | Ref. |
| 581 | *L. (L) major* | MHOM/SU/1973/5-ASKH | NA | No Amplification | Ref. |
| 561 | *L. (L) mexicana* | MHOM/BZ/1982/BEL21 | NA | No Amplification | Ref. |
| 571 | *L. (L) tropica* | MHOM/SU/1958/STRAIN-OD | NA | No Amplification | Ref. |
| 566 | *L. (V) braziliensis* | MHOM/BR/1975/M2903 | NA | No Amplification | Ref. |
| 565 | *L. (V) guyanensis* | MHOM/BR/1975/M4147 | NA | No Amplification | Ref. |
| 1365 | *L. (V) naiffi* | MDAS/BR/1979/M5533 | NA | No Amplification | Ref. |
| 1545 | *L. (V) shawi* | MCEB/BR/1984/M8408 | NA | No Amplification | Ref. |
| 1245 | *L. colombiensis* | IGOM/PA/1985/E582.34 | NA | No Amplification | Ref. |
| 3124 | *L. (L) infantum* | MHOM/PT/1989/IMT-202 | Ndel (Portugal) | Allele2/Allele2 | Strain |
| 575 | *L. (L) amazonensis* | IFLA/BR/1967/PH8 | NA | Allele2/Allele2 | Ref. |
| 563 | *L. (L) donovani* | MHOM/ET/1967/HU3 | NA | Allele2/Allele2 | Ref. |
| 2848 | *L. (L) infantum* | MCAN/BR/2005/NMT-BSB159B | Ndel (Bahia) | Allele2/Allele2 | Strain |
| 3210 | *L. (L) infantum* | MCAN/BR/2010/CALSITO I | Ndel | Allele1/Allele2 | Strain |
| 2666 | *L. (L) infantum* | MACAN/BR/2002/LVV-137 | Ndel | Allele1/Allele2 | Strain |
| 2972 | *L. (L) infantum* | MHOM/BR/2005/742EMS | Ndel | Allele1/Allele2 | Strain |
| 2668 | *L. (L) infantum* | MCAN/BR/2002/LVV-139 | Ndel | Allele1/Allele2 | Strain |
| 3685 | *L. (L) infantum* | MHOM/BR/2006/MA03-A 4P(GM) | Ndel | Allele1/Allele2 | Strain |
| 3134 | *L. (L) infantum* | MCAN/BR/2009/GRANDÃO I | Htz | Allele1/Allele2 | Strain |
| 3598 | *L. (L) infantum* | MCAN/BR/2015/TUBO9 | Del | Allele1/Allele1 | Strain |
| 3227 | *L. (L) infantum* | MCAN/BR/2010/MAGRÃO | Del | Allele1/Allele1 | Strain |
| 579 | *L. (L) infantum* | MHOM/BR/1974/PP75 | Del | Allele1/Allele1 | Strain |
| 3116 | *L. (L) infantum* | MHOM/BR/2009/HU-UFS11 | Del | Allele1/Allele1 | Strain |
| 2769 | *L. (L) infantum* | MCAN/BR/2005/SPACK | Del | Allele1/Allele1 | Strain |

The Plot (Figure 4) depicted four color-coded groups:“ no amplification” (in orange), composed by all non-*L. donovani* complex species (except for *L. amazonensis*); “Allele2/Allele2”comprising the *L. donovani* reference strain, *L. amazonensis* and two *L. infantum* from OW (Portugal and Bahia, Brazil - in blue); “Allele1/Allele2” formed by Brazilian *L. infantum* NonDEL and the HTZ sample (in green); “Allele1/Allele1” formed by all Brazilian *L. infantum* DEL (in red). The deleted site 2380 was fully amplified among OW and Brazilian NonDEL, confirming the genomic description for these samples as non-deleted. The 2330 target was fully amplified among all *L. infantum* Brazilian samples (DEL ,NonDEL and HTZ), but not for the OW strain, confirming it to be a good endogenous marker for reaction-control, possibly differentiating samples ‘origins as OW and NW – although additional OW samples should be tested. Importantly, the 2330 target allowed to distinguish between *L. donovani* complex samples and all the other assayed *Leishmania* species, except for *L. amazonensis*, also genotyped as “Allele2/Allele2” group.

**Figure 4.**
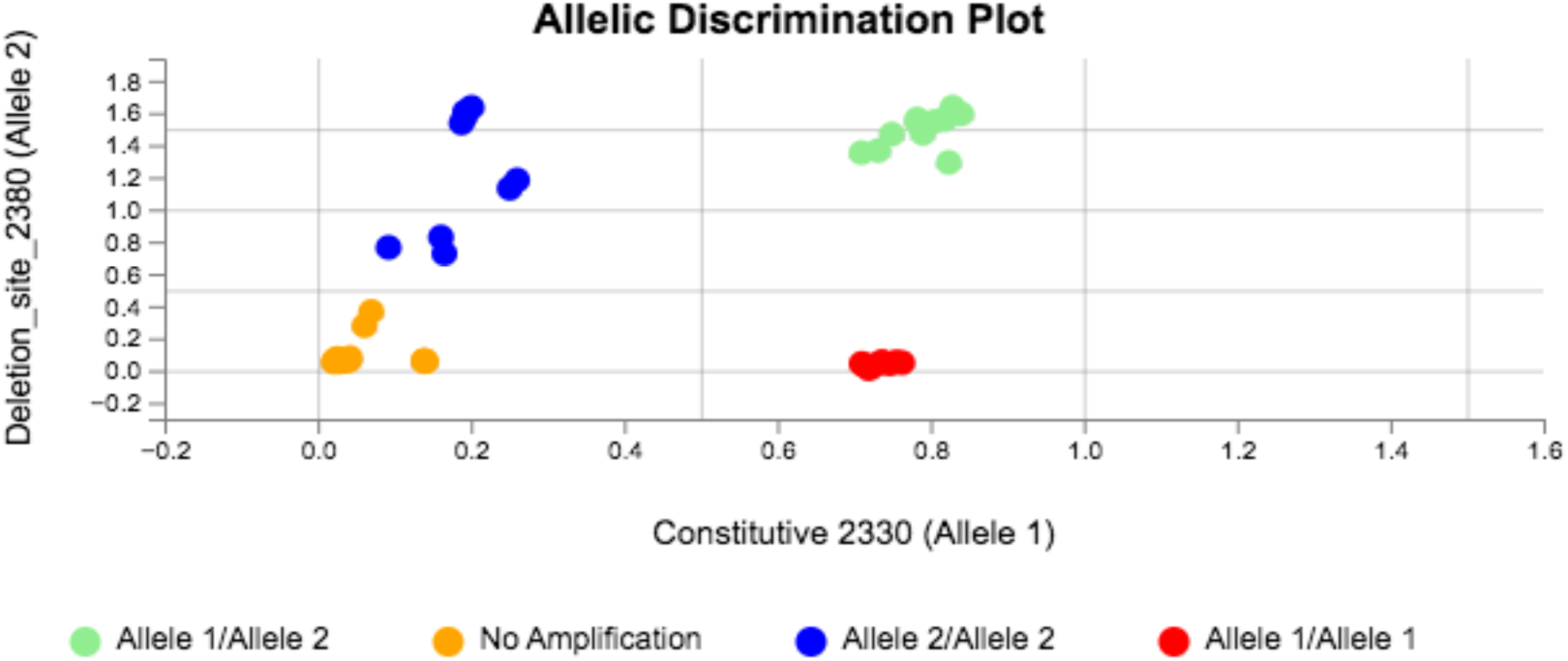
Allelic Discrimination Plot from the “Genotyping” setting applied on DNA from culture parasite from the L. (Viannia), L. (Porscisia) Paraleishmania and L. (Leishmania) subgenus. Variable samples from the L. donovani complex were specially included to represent Old World and New World Del and NoNDEL. In blue: L donovani, L. infantum from the Old World (OW) and L. amazonensis; in green: the NonDEL and HTZ New World L.; in red: the DEL samples from Brazil. The deleted site 2380 is fully amplified among blue and green samples, characterizing these strains as non-deleted; the conserved target 2330 is equally amplified for Brazilian DEL and NonDEL, confirming it as a good conserved endogenous target for genotyping New world samples. 2330 is not amplified among non-L. donovani complex samples thus representing a potential marker for differential detection for these species; moreover, 2330 is only partially amplified among L. donovani and OW L. infantum, attending thus as a potential marker for geographic origin of the sample.

Additionally, to test the protocol’s accuracy in clinical material, 28 samples collected from 18 infected dogs were included to test the approach on this type of material (Table 6). To reflect the scenario of co-circulation and coinfection by both genotypes, four samples with a mixed profile were included (MIX). All genotypes were determined beforehand by the amplification by qPCR of target 31_2380, as pointed in Figure 2.

**Table 6.**
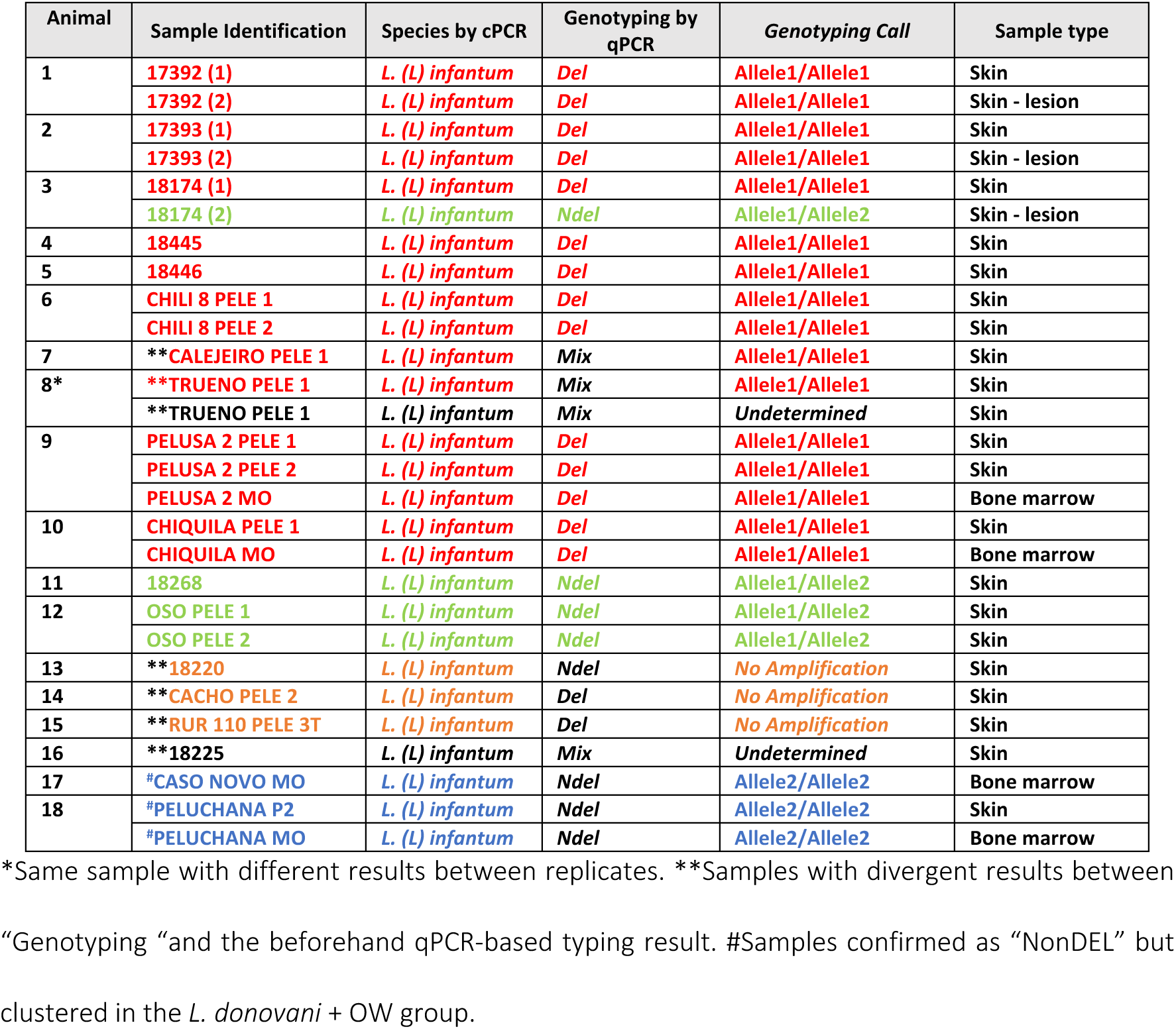
Clinical samples selected to test the differential genotyping approach by the Real time qPCR. Identification of samples, species determined by conventional PCR and sequencing, the Genotyping call (color-coded) obtained by the software and sample type.

The Allelic Discrimination Plot (Figure 5) corroborates the beforehand genotyping for 21/28 samples (72.4%). The remaining 7/28 (27.6%) samples (Table 6; marked**) had been previously characterized as “MIX” (4/7), which represents more complex samples due to the combination of genotypes, and/or presented low parasite load (Supp. Material Fig.3), explaining the conflicted outcome by the *Genotype* setting.

**Figure 5.**
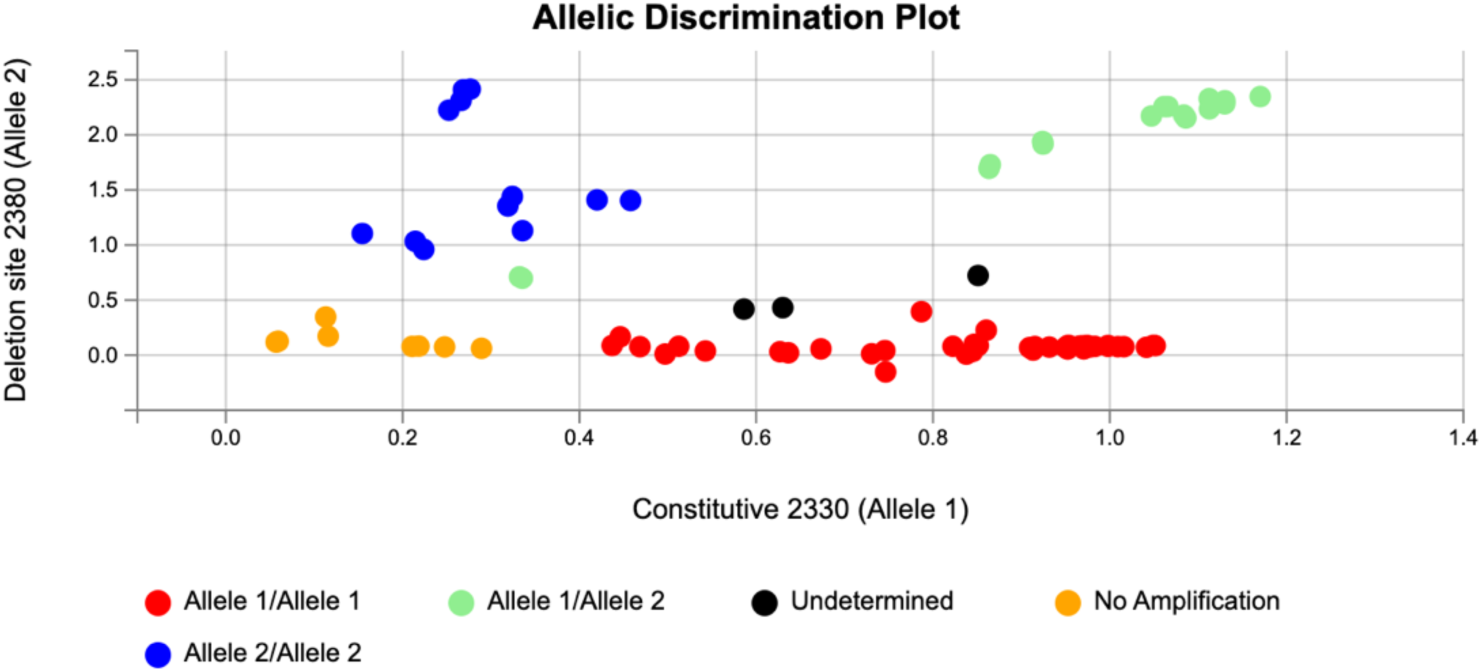
Allelic discrimination Plot for DEL, NonDEL reference samples (promastigotes) and clinical material. All clinical samples were previously confirmed as infected by L. infantum by conventional PCR for ITS1 target and sequencing. The DEL nonDEl genotype was first determined by qPCR applying absent/present amplification of targets 31.2380 and 31.2330.

### Measurement of parasite load and differential quantitation of genotypes in mixed / co-infected samples

Standard curves were obtained for 31_2330 and 31_2380 *Leishmania* targets and for the canine HPRT gene included as the host tissue equivalent (Suppl. Material Figure 1). The 28 samples from the *Genotype* setting (Table 6) and additional samples (total n=20 DEL; n=15 NonDEL; n=5 MIX) were assayed using the 31_2330 protocol, and the results were compared with those from the available highly sensitive protocol based on the kDNA target. The kDNA and 31_2330 presented a positive correlation (Pearson r=0.98; Supp. Material, Fig1)., but the mean value of equivalent of parasites/mg of tissue was higher by kDNA (1.689 for kDNA vs 0.2414 for 31_2330), as expected due to the higher number of copies for this target (Supp. Material, Table1). Blant-Altman test revealed poor agreement (Bias= -1.45, SD of Bias= 5.2) confirming that the 31_2330 protocol underestimates the parasite load.

Both protocols (31_2330 and kDNA) were applied to test the association between parasite load and the infecting genotype. No significant differences in parasite load were detected among DEL, NonDEL, and MIX groups in neither protocol (Figure 6)

**Figure 6.**
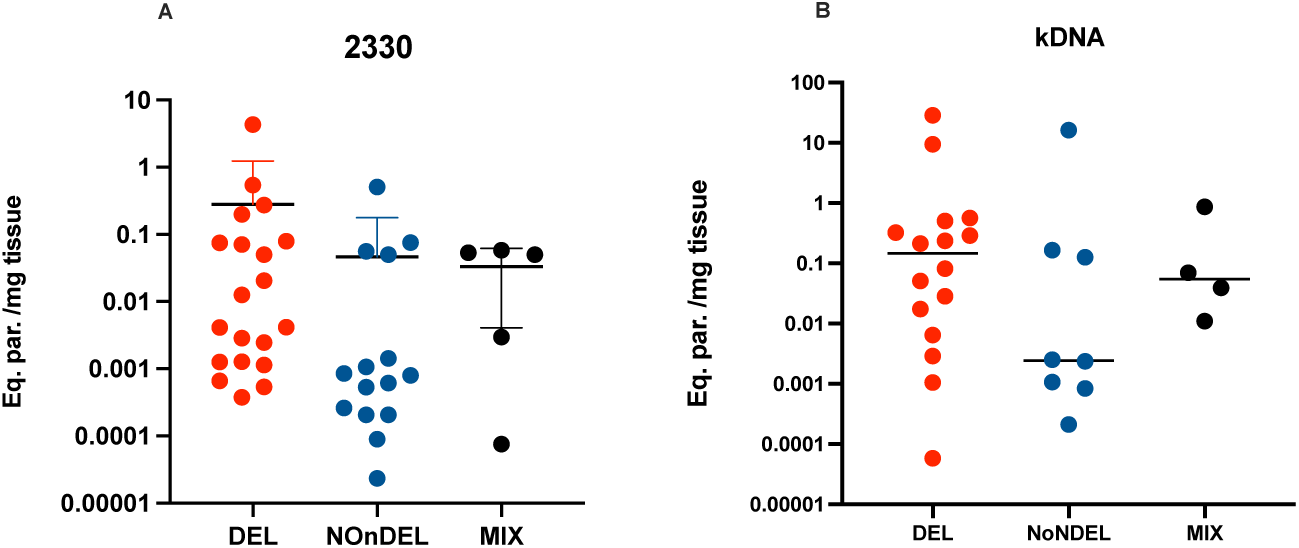
Parasite load expressed in equivalent of parasites per milligram of tissue for clinical samples infected with DEL, NonDEL or both genotypes (MIX). The values were determined by 31_2330 (n=40) and kDNA (n=28) protocols using HPRT as the host reference gene.

To be able to differentially quantify both genotypes on samples expressing the MIX profile we tested the protocol by assaying DNA from a controlled blend of DEL and NoNDEL axenic parasites. The quantity of DEL parasites was fixed while a dilution with NonDEL was prepared. The approach allowed the relatively quantification of both targets, although the absolute values by qPCR did not match accurately with the cell counting. The data shows the 31_2380 quantity fully corresponds to the serially diluted NonDEL parasites, while 31_2330 corresponds to DEL and NonDEL cells, since it is the “constitutive target” for both genotypes (Figure 7A). The limit of detection reached 0.1 NonDEL within 10^7^ DEL parasite cells.

**Figure 7.**
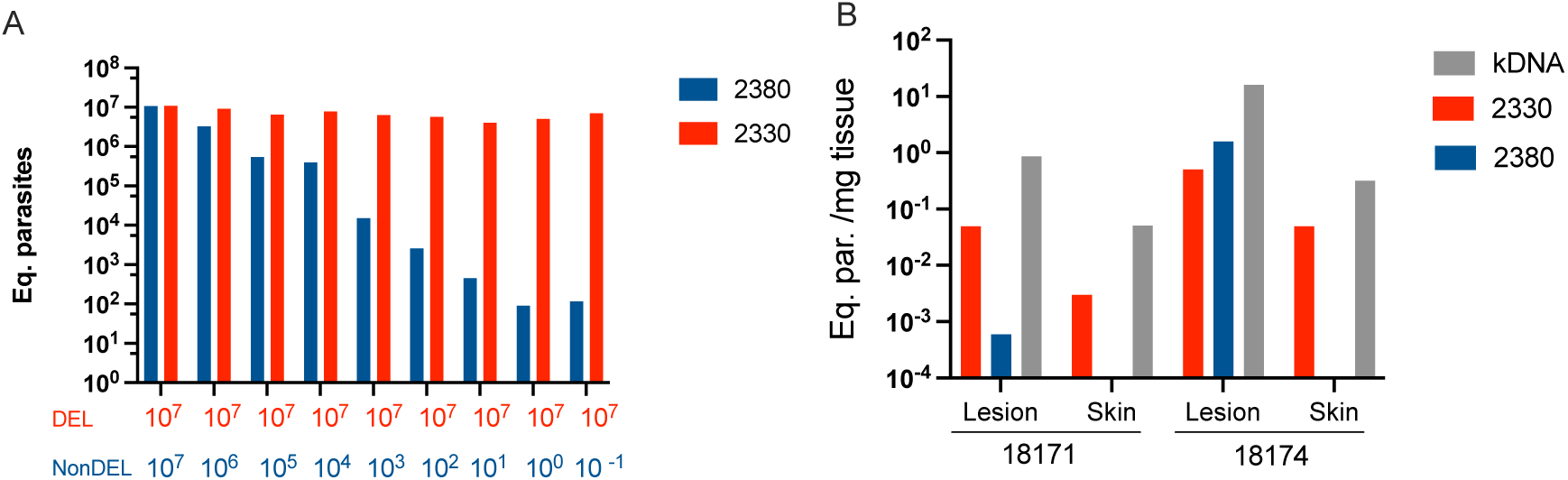
Differential quantification of 31_2330 and 31_2380 targets. A) Assay with DNA isolated from a controlled blend of DEL and NonDEL axenic parasites. The number of DEL cells was fixed while the number of NonDEL cell varied. The corresponding number of parasites (expressed as Equivalent of parasites in Y axis) obtained by the qPCR corresponds to the number of cells added to the combination by counting. Bottom; X axe: Red = number of DEL cells; Blue = number of NonDEL cells. 31_2330 detects both DEL and NonDEL parasites while 31_2380 is amplified only for NonDEL samples, allowing for the differential quantitation of both genotypes in MXI samples. B) Equivalent of parasites per mg of tissue expressed for kDNA, 31_2330 and 31_2380 from lesion and healthy skin collected from two infected dogs from Minas Gerais, Brazil. kDNA presents higher sensitivity as previously shown, but 31_2330 and 31_2380 revealed similar efficiencies and sensitivity, allowing the determination of the 31_2330/31_2380 ratio.

The approach was tested to specifically quantify the genotypes in four samples from two dogs presenting paired samples (lesion and health skin) with different genotypes (Table 7). A standard curve was prepared with the serial dilution of DNA from the skin of a healthy dog (negative for *Leishmania* infection) blended with counted NonDEL *Leishmania infantum* cells. In only in one sample (collected from 18174 skin lesion), it was possible to relatively quantify both genotypes; the additional samples presented low parasite load, outside the detection range by the standard curve.

**Table 7.** Paired samples were selected from two dogs (18171 and 18174) presenting different infecting genotypes.

| Animal | Sample | Genotype |
| --- | --- | --- |
| 18171 | (1) lesion | MIX |
|  | (2) skin | DEL |
| 18174 | (1) skin | DEL |
|  | (2) lesion | MIX |

Considering the similar efficiencies for targets 31_2330 and 31_2380 (supplemental material Fig1) the parasite loads were used to determine the 31_2330/31_2380 ratio, revealing the NonDEL genotype (detected by 31_2380) was approximately three times more prevalent (31_2330/31_2380 = 0.32) (Figure 7B).

## Discussion

The finding in the Americas of a mutant *L. infantum* carrying a 12Kb deletion (DEL) in their genome cocirculating with NonDEL strains raise epidemiological questions on the effects of these parasites in the AVL context. To answer such queries a main step is to properly detect, map and monitor the occurrence of DEL and NonDEL genotypes - the main goals of the present study.

We developed a qPCR-based strategy targeting (i) a “constitutive” genomic region, present and conserved among *L. infantum* strains, and (ii) the deleted region, which is amplified exclusively for the NonDEL, strain. The strategy proved to be accurate after being tested in DNA from cultured parasites previously characterized by WGS(17).

Then, aiming to obtain a mapping of the occurrence of genotypes, we selected a wider panel of strains deposited at CLIOC from distinct regions, including Central and South America countries. Importantly, we added to this panel DNA from clinical samples from various hosts and tissues, including those from Brazil and other South American countries. As presented before(13,17), deleted carrying *L. infantum* strains were more frequent and are widely distributed, considering both the cultured parasites and the clinical material, thus excluding the potential in vitro-selection bias of testing only isolated parasites.

Most samples were collected from dogs; thus, a fundamental question that emerges is whether the same situation would be observed for human samples. It is widely accepted that the parasite population circulating in dogs is the same as that which infects humans(20,21); also, that there is an association between canine and human cases (22,23). Both affirmations reinforce the role of dogs as major reservoirs. However, the broad range of CVL seroprevalence in Brazil (from 4 to 75 % in some regions(22)), does not usually mirror human cases. The reasons for that are yet to be fully understood, but involve complex and nonlinear heterogeneities along the transmission cycle(24). One major player might be the parasite variants. Therefore, an open question is whether DEL parasites preferentially infect the canine population or, if these parasites likewise infect humans, but lead to under-detected, asymptomatic infections and thus remain unknown. An additional concern that emerges from the higher frequency of DEL parasites among the samples from dogs regards the approved use of MIL to treat CVL. Due to the low efficacy observed during a clinical trial in Brazil, the drug was not approved to treat human visceral leishmaniasis, but despite this evidence it has been permitted to treat CVL since 2016. Studies on the effectiveness of the treatments have been conducted abroad(25) and in Brazil(15), but none of them considered whether the infecting parasite’s genotype affects the outcome. Therefore, additional studies should be pursued to determine if the current pharmacological approach is equally successful for infections by the different strains.

The MIX profile was often observed among the clinical samples assayed. In it, both genotypes are clearly present, characterizing a coinfection by both DEL and NonDEL parasites. These events are expected, as the mapping reveals the co-circulation of both genotypes, and it is thus expected coinfected hosts. Coinfection was also confirmed among paired samples from the same animal. The healthy skin and the lesion collected from two dogs and one cat revealed that, in the former only the DEL genotype was detectable, while in the sample from the lesion both genotypes (MIX) were present. The clinical and epidemiological consequence of these multiple infections are unknown and represent an additional challenge to which the current proposed protocol aims to contribute. In this regard, we applied the protocol as a differential quantitative tool, allowing for the relative quantitation of each genotype.

Although not as sensitive, the equivalent of parasites per milligram of tissue obtained by the 31_2330 qPCR positively correlated with that obtained by the kDNA- based protocol. The approach, thus, allows to concomitantly identify and quantify the infecting genotype. This tool might be especially relevant to test associations between clinical outcomes vs parasite load and infecting genotype. Having this in mind, we tested whether parasite load associates with genotypes. No difference was observed among groups. However, the samples were from very different localities, possibly from distinct transmission cycles. The same approach should thus be applied to numerically significant DEL and NonDEL samples from animals sharing the same region, preferably the same ecotype environment.

To offer the protocol as a typing tool and reduce subjectivity in genotype determination by operator evaluation—specifically through the observation of target 31_2380 amplification—and to facilitate the approach’s adoption across different laboratories, we tested the "*Genotype*" option available in the equipment software. This option, which identifies SNPs within the region amplified by primer pairs 31_2330 and 31_2380, algorithmically clusters samples and assigns genotypes based on their plot positions. This methodology groups samples through a principal component analysis format based on multiplex target amplification. Representative strains from other species and subgenera were included to assess the protocol’s potential as a differential diagnostic tool for *L. infantum* infection. The results showed high accuracy in distinguishing between DEL and NonDel samples and differentiating *L. infantum* strains from the New World (Brazil) and Old World (Europe). Reference samples representing various *Leishmania* species clustered separately from the *L. donovani* complex, highlighting "*Genotype*" as an effective differential typing tool. An exception was *L. amazonensis*, part of the *L. mexicana* complex, which clustered with the *L. donovani* complex. The reasons for this distinction in *L. amazonensis* were not identified. When applied to clinical samples, the approach also showed good accuracy, though, as occurs in other methods, it could be affected by parasite load. Interestingly, two MIX samples were classified as "undetermined," with conflicting technical replicates, indicating the method’s ability to detect complex samples containing mixtures.

## Conclusion

The qPCR-based molecular tool proposed effectively detects, identifies, and quantifies different genotypes of *Leishmania infantum*—DEL, NonDEL, and coinfections (MIX). Its application to a broad panel of clinical samples revealed the widespread occurrence of the DEL genotype, which is less susceptible to Miltefosine and harbor biological differences that may affect infection ouctomes with consequences for transmission. This highlights the need for its use in genotyping *L. infantum* infections in the Americas, especially in dogs in Brazil, where the drug is used for treatment. The protocol is thus suitable for surveillance studies and investigating associations between genotypes and relevant clinical-epidemiological phenotypes.

## Aknowledgements

This work was funded by the FIOCRUZ - PASTEUR - USP (Tripartite 2018) grant. We thank Rede de Plataformas Tecnológicas da Fiocruz-Plataforma de Análises Moleculares (RPT09J) and Coleção de Leishmania da Fiocruz (CLIOC) for providing the *Leishmania infantum* strains. We thank all the teams that participate in survaillence efforts and collection of samples, especially the Pan-America World Health Organization (PAHO-WHO). Sandro A. Pereira is recipient of productivity fellowship from CNPq, Brazil (number 316975/2023-0). We thank Heitor Herrera who coordinate the obtention of samples from coatis, in Mato Grosso do Sul. Artificial Intelligence was exclusively applied during the final stage of manuscript preparation for English language enhancement.

## Author Contributions

MCC: Investigation; Methodology; Validation; Writing – Original Draft Preparation.

ANSE: Resources; Supervision; Writing – Review & Editing.

SV: Resources; Supervision

BDC: Investigation; Methodology

CE: Investigation

AV: Resources; Writing – Review & Editing.

SAP: Resources; Supervision

GCM: Investigation; Resources

ALRR: Investigation; Resources; Supervision; Writing – Review & Editing.

MS: Investigation; Resources; Writing – Review & Editing.

CB: Resources; Supervision.

DBMF: Investigation; Resources; Supervision; Writing – Review & Editing

JMG: Investigation; Resources; Writing – Review & Editing

JET: Resources; Supervision; Writing – Review & Editing

LFB: Investigation; Resources; Writing – Review & Editing

ML: Resources; Supervision; Writing – Review & Editing

LPV: Resources; Investigation; Methodology; Writing – Review & Editing

CR: Resources; Supervision; Writing – Review & Editing.

OCM: Resources; Methodology; Writing – Review & Editing.

EC: Conceptualization; Data Curation; Resources; Supervision; Writing – Review & Editing.

MCB: Conceptualization; Data Curation; Formal Analysis; Funding Acquisition; Methodology; Project Administration; Resources; Supervision; Visualization; Writing – Review & Editing.

